# Air and surface measurements of SARS-CoV-2 inside a bus during normal operation

**DOI:** 10.1101/2020.06.26.173146

**Authors:** Piero Di Carlo, Piero Chiacchiaretta, Bruna Sinjari, Eleonora Aruffo, Liborio Stuppia, Vincenzo De Laurenzi, Pamela Di Tomo, Letizia Pelusi, Francesca Potenza, Angelo Veronese, Jacopo Vecchiet, Katia Falasca, Claudio Ucciferri

## Abstract

Transmission pathways of SARS-CoV-2 are through aerosol, droplet and touching infected material. Indoor locations are more likely environments for the diffusion of the virus contagion among people, but direct detection of SARS-CoV-2 in air or on surfaces is quite sparse, especially regarding public transport. In fact, an important demand is to know how and if it is safe to use them. To understand the possible spreading of COVID-19 inside a city bus during normal operation and the effectiveness of the protective measures adopted for transportation, we analysed the air and the surfaces most usually touched by passengers. The measurements were carried out across the last week of the lockdown and the first week when gradually all the travel restrictions were removed.

## Introduction

The spread of severe acute respiratory syndrome coronavirus 2 (SARS-CoV-2) affected over 100 countries in a matter of weeks, therefore on January 30th the World Health Organization (WHO) announced the COVID-19 epidemic as a Public Health Emergency of International Concern [1]. Italy has been one of the most affected countries with more than 230000 infected people and more that 33000 deaths and became the first country in Europe to proceed with a total lockdown (so called phase 1, started on 9 March 2020). The government decided to impose strong restrictions in the whole Country closing schools, public places (such as, restaurants or cafés) and shops, allowing only the basic necessities stores (such as supermarkets and pharmacies) and relative activities to remain open [2]. The huge increase number of infected people resulted, on 13 March, in more striking measures including the transport rationalization with a strong reduction of public transport, maintaining a minimum level of services [3]. A protocol between the Italian Ministry of Infrastructure and Transport and the Italian Ministry of Health, together with trade organizations and trade union representatives, established anti-contagion rules and actions and promoted cleaning and disinfection procedures of public transport services to contain the COVID-19 spreading. Moreover, it was performed with the aim to ensure the safety of workers and travellers in the transport and logistics sectors [4]. One of the main measures recommended was the recurrent cleaning and disinfection of frequently touched surfaces such as handles and rails because of the potential environmental stability of SARS-CoV-2 that, according to some reports, could span from up to three hours in the air post-aerosolisation to about 24 hours on cardboard and about three days on plastic and stainless steel [5]. Recent studies shown the possible airborne transmission of the virus in public places, which could be spread by asymptomatic people [6, 7]. In addition, the research findings suggest reducing the number of people in the same ambient and carry out control actions to limit the pandemic expansion [6]. Therefore, for buses and trains sanitation were recommended to be performed with virucidal licensed products, based on sodium hypochlorite, or those based on ethanol (at least 70%), after cleaning with a neutral detergent [8]. In Italy, the end of the lockdown was planned to finish gradually with different dates for the reduction of constrains as function of the infection risks of the different activities. Starting on 18 May 2020, phase 2 began with a Decree of the President of the Council of Ministers [9] that included guidelines for public transport establishing general rules, such us: 1) reduction of the number of passengers inside the buses, 2) interpersonal distance of one meter, 3) rear door boarding in order to protect drivers, 4) only distanced seats permitted, 5) passengers must frequently sanitize hands and 6) obligation to wear facial masks [9]. The local governments, following the national guidelines from the DPCM [9], established the exact operational rules for local public transportation system. In details, the Abruzzo Region (Central Italy), where this study was carried out, defined that: 1) the maximum number of passengers on-board buses must not exceed 40% of the total seats and 15% of standing places, if provided; 2) standing places must be marked with a signal on the ground 3) by 18 May 2020 at least 50% of the services performed before the reduction due to COVID-19 is reactivated, reaching the 70% within and not beyond 31 May 2020 [10].

Analysis of the air and surfaces in indoor environments are crucial to better understand the SARS-CoV-2 spreading and airborne transmission, to better assess the risks for doctors and health-care operators [6]. However, these observations are very limited and mostly confined to hospital environments [11, 12]. Results of risk models assessing the airborne transmission of the virus in different indoor environments such as restaurants, post offices, pharmacies, supermarkets and banks, suggest the key role of air ventilation, but simulations of more confined environments like those inside city buses, trolleybuses, trams or trains, are missing [13]. For this reason, we analysed both the air and samples taken from the surfaces most frequently touched of a city bus, to better understand the possible spreading of SARS-CoV-2 and in order to assess the effectiveness of the measures defined for the containment of COVID-19 diffusion.

## Materials and methods

The study was conducted from 12 to 22 May 2020 in Chieti, a town in the Abruzzo region that is the fifth Italian region for mortality due to COVID-19, with an infection fatality rate (deaths / cases) of 12.1% [14]. In Abruzzo, as of 28 May 2020, are reported 3237 cases of infected people, 820 in the Chieti province, which is 0.213% of the total population [15]. In the present study, the environmental inside the trolleybus line number 1 of the local transportation systems was monitored. This line is the most important of the town in terms of number of passengers, covering a route of 20 km with 50 stops that from downtown reaches the University Campus and the Santa Annunziata Hospital and then back to downtown.

The samples of air inside the bus were carried out every day of the two observational weeks, excluding weekends, during one shift (5 routes) of the line 1 that started at 12.00 and finished at 18:30. The bus was operated following the rules established by the DPCM [9] and the DPRC [10]. Two microbiological gelatine membrane sample filters of 80 mm diameter were installed: one close to the ticket machine, the other on the rear part of the bus (Fig 1). These filters are the proper support, to be analysed with the RT-PCR, for the detection of SARS-CoV-2 virus [11]. In fact, the microbiological gelatine membrane filters, employed in this study, were tested at the Clinic of Infectious Diseases of the S.S. Annunziata Hospital in Chieti to check their performance in detecting the virus in air. It has been found, in some cases, that samples collected in an isolation room with patients symptomatic to the SARS-CoV-2 were positive to the virus.

**Fig 1.**
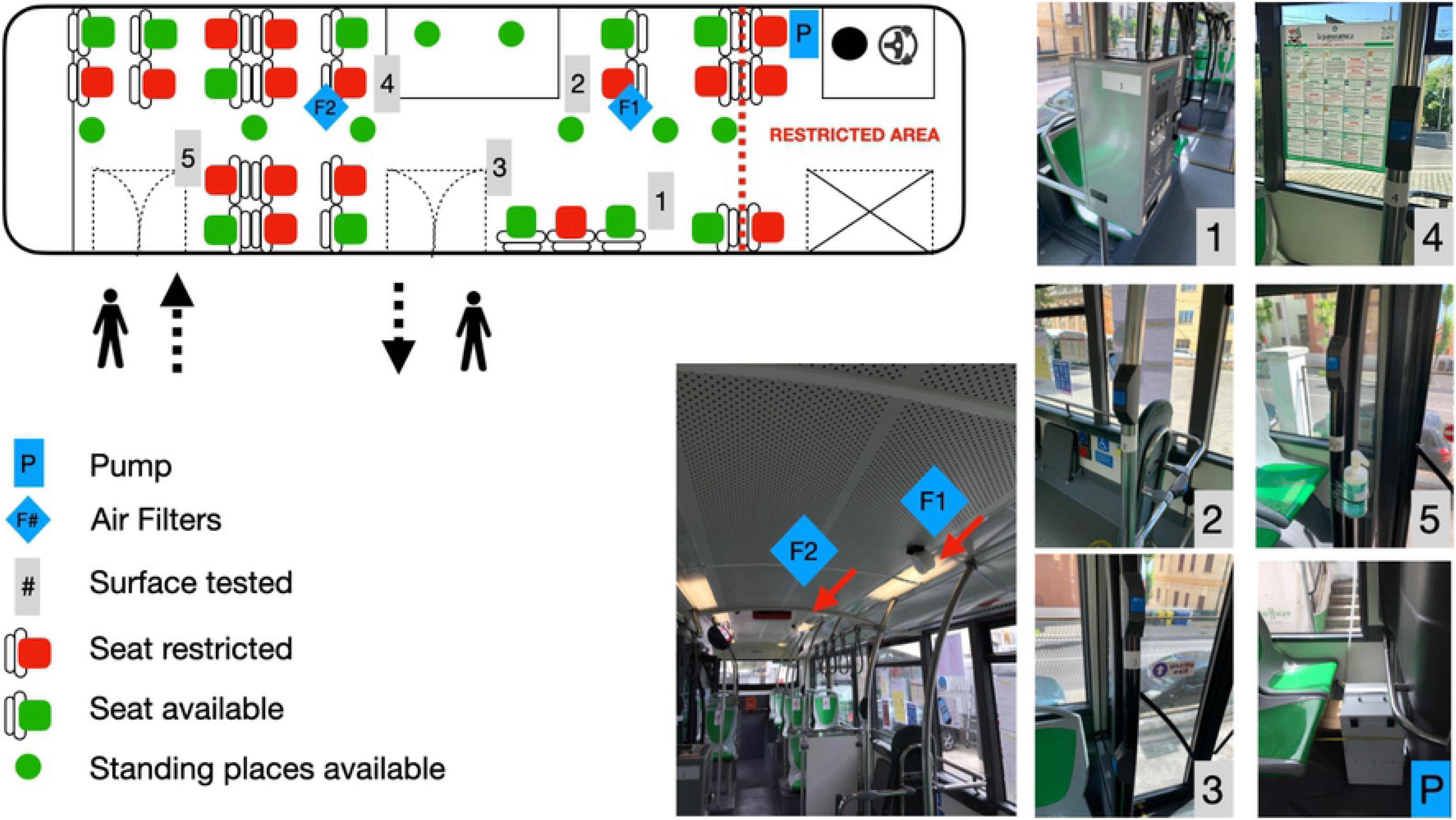
Sketch of the trolleybus where samples were carried out. On the left there is a schematic of the inside the trolleybus showing the restrictions in terms of seats and standing places, to follow the protocol to reduce the risk of transmission of the COVID-19 virus. On the right, are shown pictures and places of the surfaces where samples were taken and the position of two filters for air analysis. The number of the pictures correspond to the sample points reported in the trolleybus drawing.

A pump system, fed by the trolleybus electrical power supply, ensured an air flow of 24 l/min to each filter. All the air samples were carried out during the 6.5 hours daily operation of the bus. One air sample, as a control reference, was taken without passengers overnight, for 21 hours, with the bus in the hangar.

Surface samples were carried out with wet swabs on five points on the bus that are those more frequently touched by the passengers, according to the experiences of the bus drivers (Fig 1). The samples for each observation day and each surface were taken before the beginning of the bus shift, to have a reference, and immediately after the end of the shift. Cleaning and sanitation inside the bus are carried out daily using sodium hypochlorite 0.1% and ethanol 70%. Once a week the bus is further sanitised with an electric aerosol applicator that delivers for 1 minute, highly oxidizing, non-foaming acid disinfectant, containing 56 g/l of peracetic acid, 12 g/l of hydrogen peroxide and 56 g/l of acetic acid. Moreover, once a week, the entire trolleybus cabin is ozonized for 10 minutes.

Surface and air samples were collected and delivered immediately after gathering them to the Microbiology and Molecular Genetics laboratory of the Center for Advanced Studies and Technology (CAST), University “G. d’Annunzio” of Chieti-Pescara to be analyzed through RT-PCR technique. The collected samples (wet swabs and microbiological gelatine membrane) were inserted on 2 cc of physiologic solution and transported to the laboratory. On arrival at the research lab, specific real-time reverse transcriptase-polymerase chain reaction (RT-PCR) (TaqMan™ 2019-nCoV Assay Kit v2; Thermo Fisher Scientific, Italy) targeting RNA-dependent RNA polymerase was used to detect the presence of SARS-CoV-2 [16, 17]. This technique uses 3 genes: ORF1ab, N gene and S gene to quantify the viral load with a number of cycles for the fluorescent signal to cross the threshold in RT-PCR. The threshold is 5000, baseline is 5 and cut-off is 37 cycles. Lower values of the number of cycles means higher viral load. Samples are considered “Positive” when at least two genes have a cycle threshold value < 37; if cycle threshold value is ‘Undetermined’ or >37 for two or all the genes, then the result of the sample is “Negative”.

## Results and discussion

During the whole observation period about 1100 passengers travelled using the trolleybus set up for the observations reported here, with an average of 123 people for each measurement shift as shown in details in Table 1. All the surfaces samples were negative for two or all the genes to RT-PCR analysis for SARS-CoV-2 virus (‘undetermined’ or >37). Similarly, the same results were obtained for all the air samples during the whole study period.

**Table 1.** Overview of the observations performed inside the trolleybus, where the columns ‘Before the bus shit’ are the surfaces sampled before the beginning of bus journey, ‘After the bus shift’, are the surfaces sampled after end of the bus journey. Air samples were carried out on two sample sites (F1 and F2, see Fig 1) every day, excluding the first two observational days. The values reported (>37) for all the samples means that they are ‘negative’ to the presence of the SARS-COV-2 virus since none of them showed at least two of the three genes (ORF1ab, N gene and S gene) with a cycle threshold value < 37 (see the text).

|  |  | Before the bus shift |  |  |  |  | After the bus shift |  |  |  |  | Air Sample |  | Passengers |
| --- | --- | --- | --- | --- | --- | --- | --- | --- | --- | --- | --- | --- | --- | --- |
| Sample point |  | 1 | 2 | 3 | 4 | 5 | 1 | 2 | 3 | 4 | 5 | F1 | F2 |  |
| Tuesday | 12/05/2020 | >37 | >37 | >37 | >37 | >37 | >37 | >37 | >37 | >37 | >37 | - | - | 202 |
| Wednesday | 13/05/2020 | >37 | >37 | >37 | >37 | >37 | >37 | >37 | >37 | >37 | >37 | - | - | 60 |
| Thursday | 14/05/2020 | >37 | >37 | >37 | >37 | >37 | >37 | >37 | >37 | >37 | >37 | >37 | >37 | 75 |
| Friday | 15/05/2020 | >37 | >37 | >37 | >37 | >37 | >37 | >37 | >37 | >37 | >37 | >37 | >37 | 160 |
| Monday | 18/05/2020 | >37 | >37 | >37 | >37 | >37 | >37 | >37 | >37 | >37 | >37 | >37 | >37 | 109 |
| Tuesday | 19/05/2020 | >37 | >37 | >37 | >37 | >37 | >37 | >37 | >37 | >37 | >37 | >37 | >37 | 106 |
| Wednesday | 20/05/2020 | >37 | >37 | >37 | >37 | >37 | >37 | >37 | >37 | >37 | >37 | >37 | >37 | 116 |
| Thursday | 21/05/2020 | >37 | >37 | >37 | >37 | >37 | >37 | >37 | >37 | >37 | >37 | >37 | >37 | 141 |
| Friday | 22/05/2020 | >37 | >37 | >37 | >37 | >37 | >37 | >37 | >37 | >37 | >37 | >37 | >37 | 138 |
| Total passengers |  |  |  |  |  |  |  |  |  |  |  |  |  | 1107 |

These results mean that none of the samples resulted ‘Positive’ to the SARS-CoV-2 virus, on the bus surfaces and indoor air. Unfortunately, we could not test each passenger for SARS-CoV-2, therefore we do not know exactly how many infected people travelled on the trolleybus during our observations. A recent work, based on the analyses of data from different parts of the world (China, Italy, US, Greece) and diverse situations, suggests that the asymptomatic people infected by SARS-CoV-2 are between 40% and 45% of the population [18]. Considering a conservative estimation of 30% asymptomatic people infected by SARS-CoV-2, since 123 passengers travelled on average for each bus shift, we estimated that about 37 infected and asymptomatic people potentially touched the surfaces that we sampled at the end of the journeys and breathed inside the bus while our instrument was sampling the indoor air. Under this hypothesis we can argue that the requirements of wearing gloves and cleaning up hands, using a dispenser of alcohol-based sanitizer at the bus entrance door, seem to keep the surfaces and the air inside the bus safe and free from SARS-CoV-2 virus. At the same time the rules of wearing a facial mask during travelling, and the recommendation to keep the windows open during bus riding to allow high air ventilation, probably prevent the virus diffusion in the air inside the bus. These results are in agreement with different model simulations that recommend facial masks to combat the SARS-CoV-2 virus spread in aerosols and droplets by asymptomatic people [19]. Moreover, the air ventilation, that model simulations showed to be important to reduce the risk of virus transmission in different indoor environments [13], is confirmed to be essential also in a more confined location like inside a bus.

## Conclusion

The end of the lockdown imposed to contain the COVID-19 infection outbreak is entailing a growing number of people that restart the usual daily activities including travelling on public transport. Our observations inside a bus showed that the air and all the surfaces samples were not infected by SARS-CoV-2 virus. Even if it was not possible to test the passengers to SARS-CoV-2 but considering that the asymptomatic people infected could be more than 30% [18], we can expect a potential infection inside the bus. Whether or not the number of infected passengers was about 30%, our findings confirm that the measures established for public transport in terms of sanitation, air ventilation and interpersonal precautions (facial mask, distancing, hands hygienisation) are effective, at least during this study, to make healthy and COVID-free the environment inside the buses.

## Author contributions

**Conceptualization:** Piero Di Carlo, Piero Chiacchiaretta, Bruna Sinjari, Eleonora Aruffo

**Methodology:** Piero Di Carlo, Piero Chiacchiaretta, Bruna Sinjari, Eleonora Aruffo

**Bus measurements:** Piero Di Carlo, Piero Chiacchiaretta

**Hospital measurements:** Jacopo Vecchiet, Katia Falasca, Claudio Ucciferri

**RT-PCR analysis and data validation:** Liborio Stuppia, Vincenzo De Laurenzi, Pamela Di Tomo, Letizia Pelusi, Francesca Potenza, Angelo Veronese

**Writing – original draft:** Piero Di Carlo, Piero Chiacchiaretta, Bruna Sinjari, Eleonora Aruffo

**Writing – review & editing:** Piero Di Carlo, Piero Chiacchiaretta, Bruna Sinjari, Eleonora Aruffo, Liborio Stuppia, Jacopo Vecchiet

## Acknowledgments

We want to thank the staff of the La Panoramica Group and the Municipality of Chieti for kindly providing the trolleybus, the technical support, and everything needed to complete this scientific work. We thank Mrs Elvira D’Annunzio, Mrs Maria Di Biase and Mrs Daniela Romano for their administrative support. We thank Mrs Manuela Rastelli for the revision and improvement of English. This work was partially supported by DISPUTER research funds (ex-60%).

